# Reassessing the Role of Foot Power in Human Gait

**DOI:** 10.1101/2023.10.11.561917

**Authors:** Quinn Yetman, Lauren Welte, Aidan Shimizu, Michael J Rainbow

**Affiliations:** Department of Mechanical and Materials Engineering, Queen’s University, Kingston, Canada; Mechanical Engineering, University of Wisconsin – Madison, Madison, United States

**Keywords:** foot power, arch spring, upright gait, biplanar videoradiography, walking, running

## Abstract

The foot acts as the primary interface to the ground during bipedal locomotion. It absorbs and returns energy over stance as the longitudinal arch deforms and recoils. The term ‘arch recoil’ evokes the concept that the foot’s returned energy directly propels the centre of mass forward by lifting the talus. However, recent work has shown that arch recoil does not directly drive the body forward; instead, it lowers and posteriorly tilts the talus, putting it into a more favourable position for upright gait. Here, we aim to supply a kinetic explanation for this mechanism. We applied the unified deformable power approach to highly accurate talus kinematics from biplanar videoradiography and force plate measurements to measure the power absorbed/produced by the foot. We coupled these measurements with a simple mathematical model that allowed us to restrict rotation and linear actuation of the talus caused by the recoil of the arch to demonstrate that positive foot power primarily contributes to posteriorly tilting the talus. This suggests the role of positive foot power during propulsion is to keep the talocrural surface in a more favourable position for upright gait rather than directly propelling the centre of mass forwards. These findings highlight that arch mobility during push-off is critical for allowing the ankle to directly propel the body forward and upward during the propulsive phase of gait.

## Introduction

The foot has mobile arches below the talocrural or true ankle joint, enabling it to manage our mechanical interactions with ground. The foot’s function is important, for example when its mobility is limited the metabolic cost of running increases^1^. The tissues spanning the longitudinal arches absorb power during the first half of stance, as the arches flatten, and generate power during push-off, as the arches recoil^2–15^. The timing of positive power production implies that arch recoil directly propels the body’s center of mass during locomotion by lifting the talus forward and upward^1,2,16^; however, a recent study that focused on the kinematics of the longitudinal arch argued that if the foot remained rigid, the body’s centre of mass would be higher and more forward compared to when the arch recoiled (Figure 1a)^17^. If positive propulsive foot power due to arch recoil does not serve to directly propel the centre of mass by driving the talus forward and upward, what alternative function does it serve?

**Figure 1:**
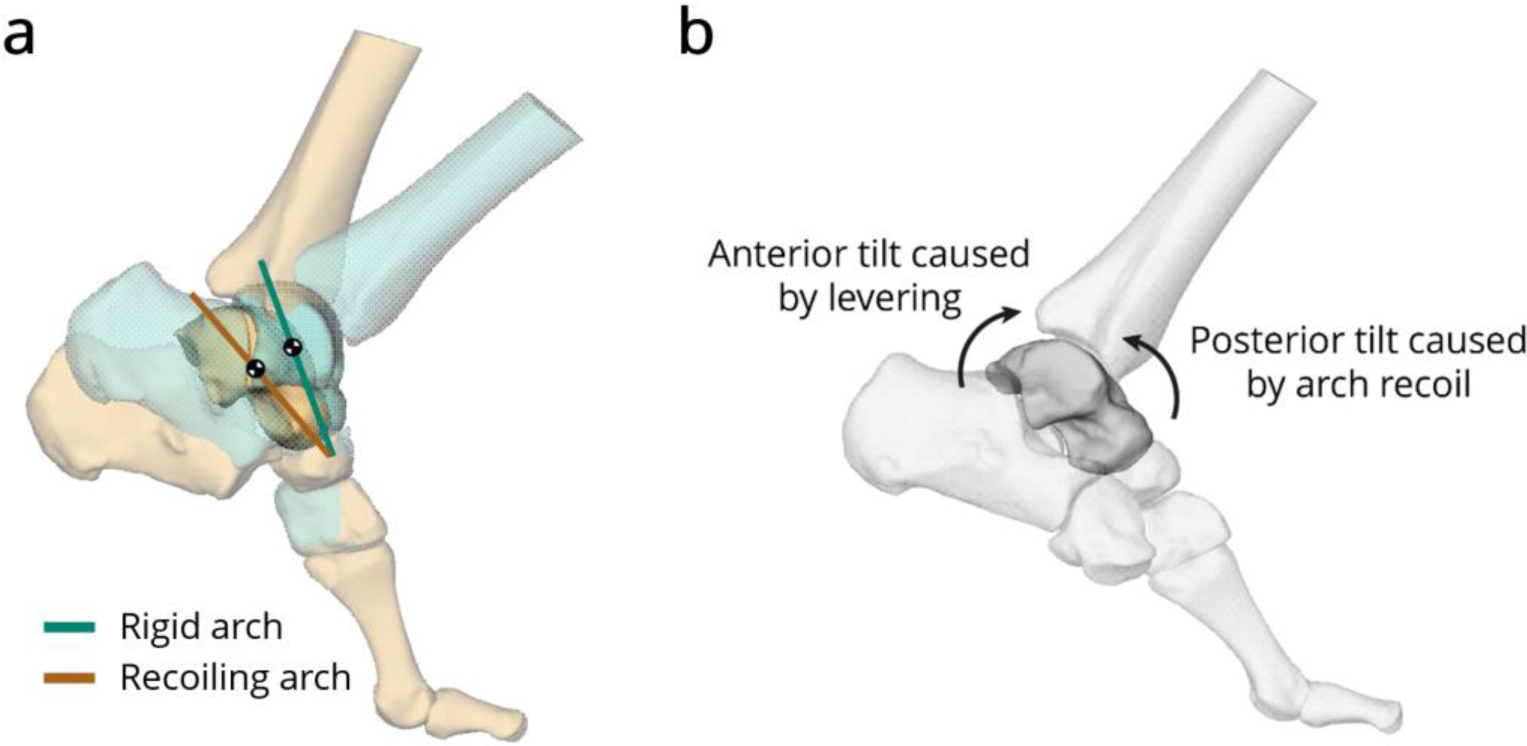
(a) Arch deformation causes the talus to rotate backwards on the arch, keeping the talocrural surface level and putting the centre of mass in a more inferior and posterior location than if the arch were rigid. (b) The recoil of the arch introduces posterior tilt to the talus in the global frame while the levering of the foot about the metatarsal heads causes anterior tilt in the global frame.

In what we call the “arch recoil-upright gait hypothesis”, arch recoil during propulsion supports ankle function by keeping the talocrural surface level, allowing the tibia (and the body) to remain more upright^17^. As the arch recoils, it tilts the talus posteriorly relative to the first metatarsal (Figure 1a). Looking from a global reference frame, this means that while the foot levers about the metatarsal heads and the talus tilts anteriorly in global, arch recoil rotates the talus in the opposite direction – effectively subtracting some global talus anterior tilt and lowering the body’s centre of mass (Figure 1b). Therefore, the “arch recoil-upright gait hypothesis” predicts that positive arch recoil power acts to posteriorly tilt the talus, effectively slowing its global anterior tilt. This is counterintuitive because positive power occurs when the velocity and angular velocity are in the same direction as the force and moment. In contrast, the upright gait hypothesis predicts that positive power slows the global talus anterior tilt that is caused by ankle plantar flexion.

In this paper, we seek to provide theoretical and experimental evidence to test our prediction that positive foot power is used to tilt the talus posteriorly relative to how it would tilt if the foot remained rigid during push-off. The unified deformable (UD) power approach is ideally suited for testing our prediction because it simplifies the foot’s complexity by assuming all structures distal to a proximal rigid segment are massless and deformable^18,19^. When the talus is used as the proximal rigid segment for the UD power approach, it captures foot power. It is useful for understanding how the foot generates power during recoil because it treats the talus in the global reference frame, allowing us to examine how adding posterior tilt from the arch to the anteriorly tilting talus (from ankle plantarflexion) affects arch power. An important caveat here is that the foot’s arch also shortens as it recoils^17,20–22^, and it is not obvious how to decouple the contributions to power from arch rotation versus arch shortening. A mathematical model that is guided by in vivo measurements of arch recoil and ground reaction forces may allow us to determine the contributions of arch shortening and rotation to overall foot power.

Here, we test the prediction that positive foot power during propulsion is used to posteriorly tilt the globally anteriorly tilting talus. We applied the UD power model to highly accurate biplanar videoradiography (BVR) measurements of the talus in motion integrated with ground reaction forces during walking and running. To help illustrate how arch recoil generates positive net power relative to a rigid arch that produces no power during ankle plantar flexion, we decomposed the UD power into its rotational (moment * angular velocity) and translational (force * linear velocity) components. We then coupled our in vivo measurements to a simple mathematical model that allowed us to understand the contributions of shortening and posterior tilt on arch recoil power. We also examined kinetic timing of foot and ankle power to test whether they are coupled, lending further support to our prediction that arch recoil assists ankle function. Finally, we place our values in context by computing the positive, negative, and net work done by the foot, ankle-foot, and ankle complexes during walking and running, enabling researchers without access to the talus to determine how computing foot power with the commonly used calcaneus as the proximal rigid segment differs.

## Results

### Foot, Ankle-Foot, and Talocrural Power

As expected, the foot, ankle-foot, and talocrural joint absorb power in the first part of stance then generate power later in stance for both walking and running (Figure 2). To determine how the distal structures rotate and translate the talus, we further separated the foot power (talus as proximal rigid segment) into its rotational and translational components (Figure 3). Rotational power derives from the deformable structures applying moments to rotate the talus while translational power derives from the deformable structures applying forces to translate the talus. During propulsion, we found the translational power had a large positive peak while the rotational power had a large negative peak. When added together this gave a net foot power during propulsion that started negative before becoming positive as the translational component became larger in magnitude than the rotational component.

**Figure 2:**
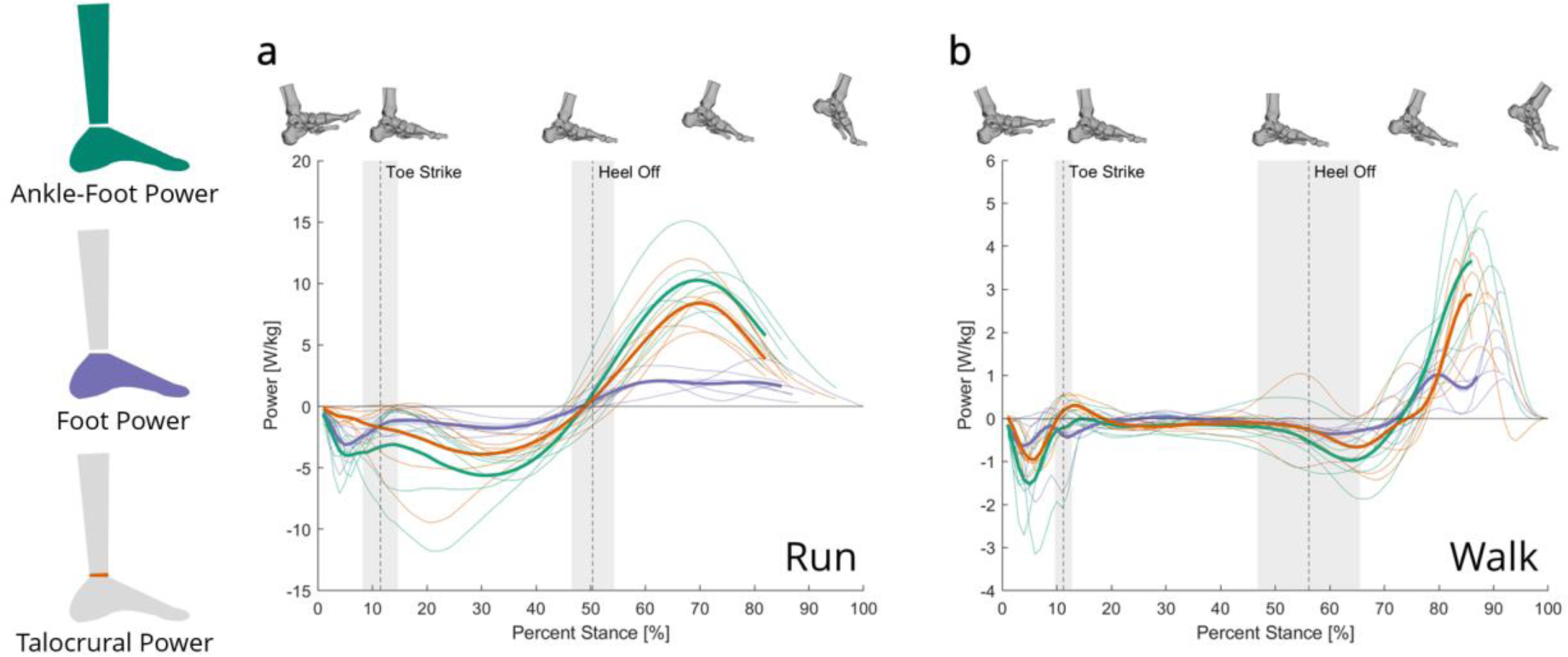
The unified deformable (UD) power over stance calculated using the tibia (ankle-foot power) and the talus (foot power) as the proximal rigid segment for (a) running and (b) walking. The talocrural power is calculated by subtracting the foot power from the ankle-foot power. Thin lines represent each participant (n = 6), and the thick lines represent the mean (mean lines stop when data becomes unavailable for a single participant due to limitations with the BVR capture volume). Mean ± standard deviation of toe strike and heel off are indicated with dashed lines and gray shading.

**Figure 3:**
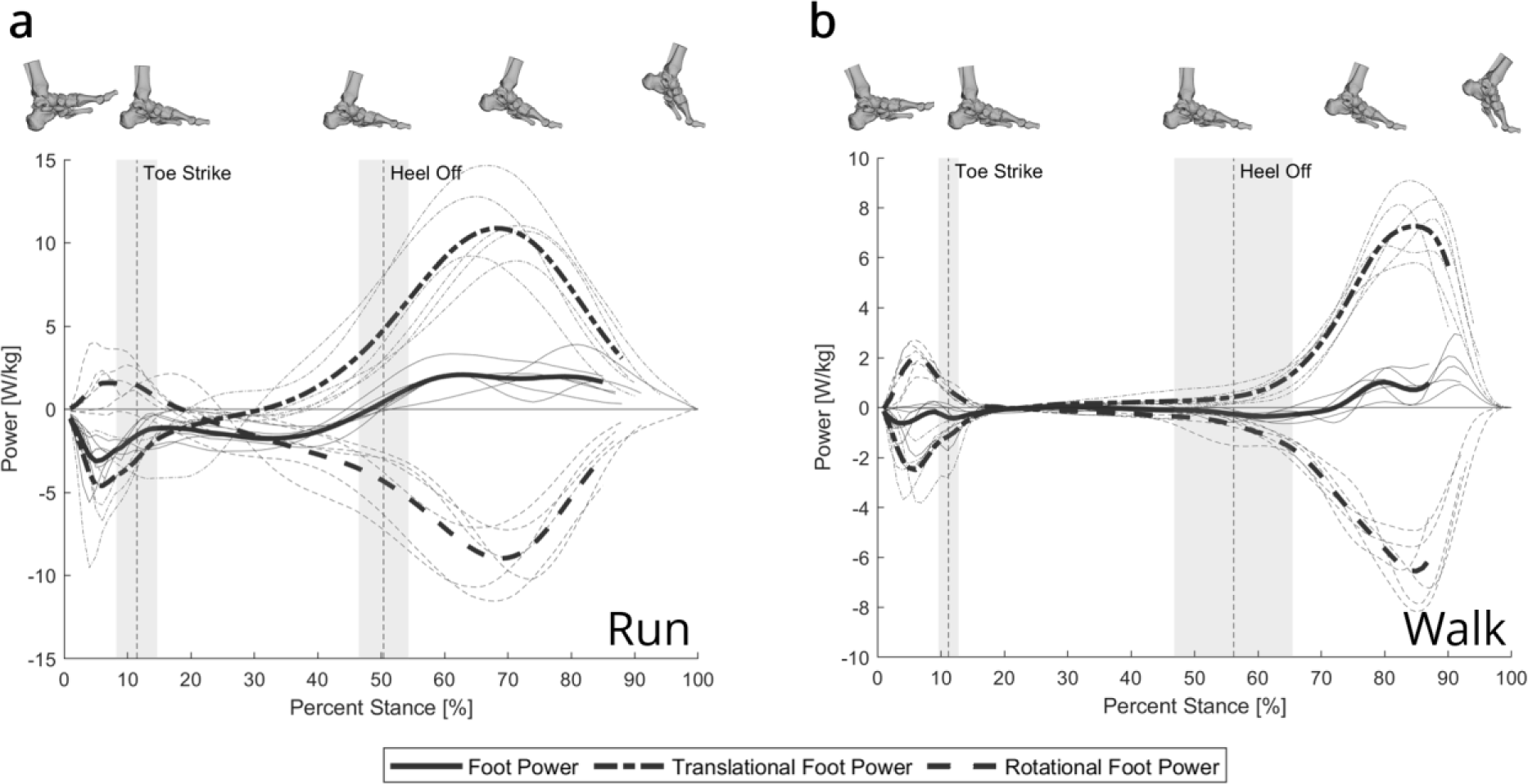
The unified deformable (UD) foot power calculated using the talus and its rotational and translational components over stance for (a) running and (b) walking. Thin lines represent each participant (n = 6), and the thick lines represent the mean (mean lines stop when data becomes unavailable for a single participant). Mean ± standard deviation of toe strike and heel off are indicated with dashed lines and gray shading.

### Simple Foot Model

To provide insight into how arch recoil impacted the rotational and translational components of the foot power such that the foot begins to generate positive power, we developed a simple, two-dimensional model of the foot, composed of a mass that rotates on the end of a massless lever arm with changing length (Figure 4). The mass is analogous to the talus, and the fulcrum of the lever arm is analogous to the metatarsal heads. The rotation of the lever arm about the fulcrum represents the foot levering about the metatarsal heads as the ankle plantarflexes, while the additional rotation of the mass relative to the lever arm, deemed “recoil rotation”, represents the rotation of the talus caused by arch recoil. We also captured lever arm shortening, deemed “recoil shortening”. We did this because the distance between the talus and first metatarsal decreases while the arch recoils. We fit model parameters to a walking trial and calculated the UD power using the mass as the proximal rigid segment (the experimentally driven case), which allowed us to test how arch recoil shortening and arch recoil rotation independently affect foot power.

**Figure 4:**
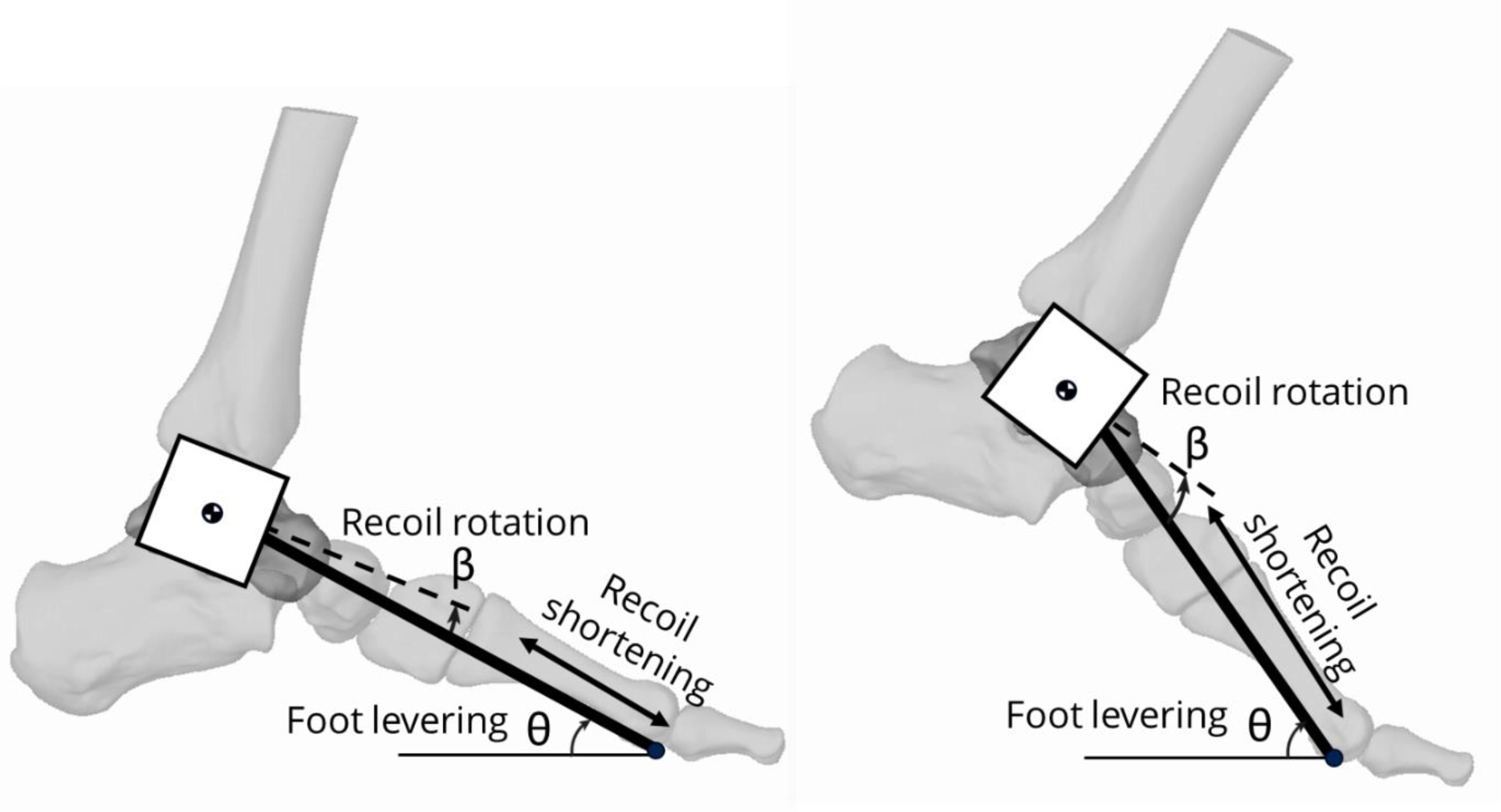
The simple lever model that represents the foot during propulsion. A mass at the end of a massless lever can rotate (representing the talus rotating) and shorten (representing the talus translating toward the metatarsal head) as it rotates about a pivot point.

We started with the rigid lever case (no recoil rotation or shortening) to see how a completely rigid arch affected rotational, translational, and net foot power (Figure 5a). As expected, we found the foot produced no net power during propulsion as there were no deformable elements. The rotational and translational power are nonzero but perfectly offset one another. This is because the translational power terms (F_GRF_, v) are in the same direction while the rotational terms (M_GRF_, ω) are in the opposite direction (Figure 6). Since the distance between the centre of mass of the talus and the ground is constant (r_COP_), the magnitude of the rotational component of power exactly equals the magnitude of the translational component of power (see Supplemental S1 for worked out equations), resulting in zero net power.

**Figure 5:**
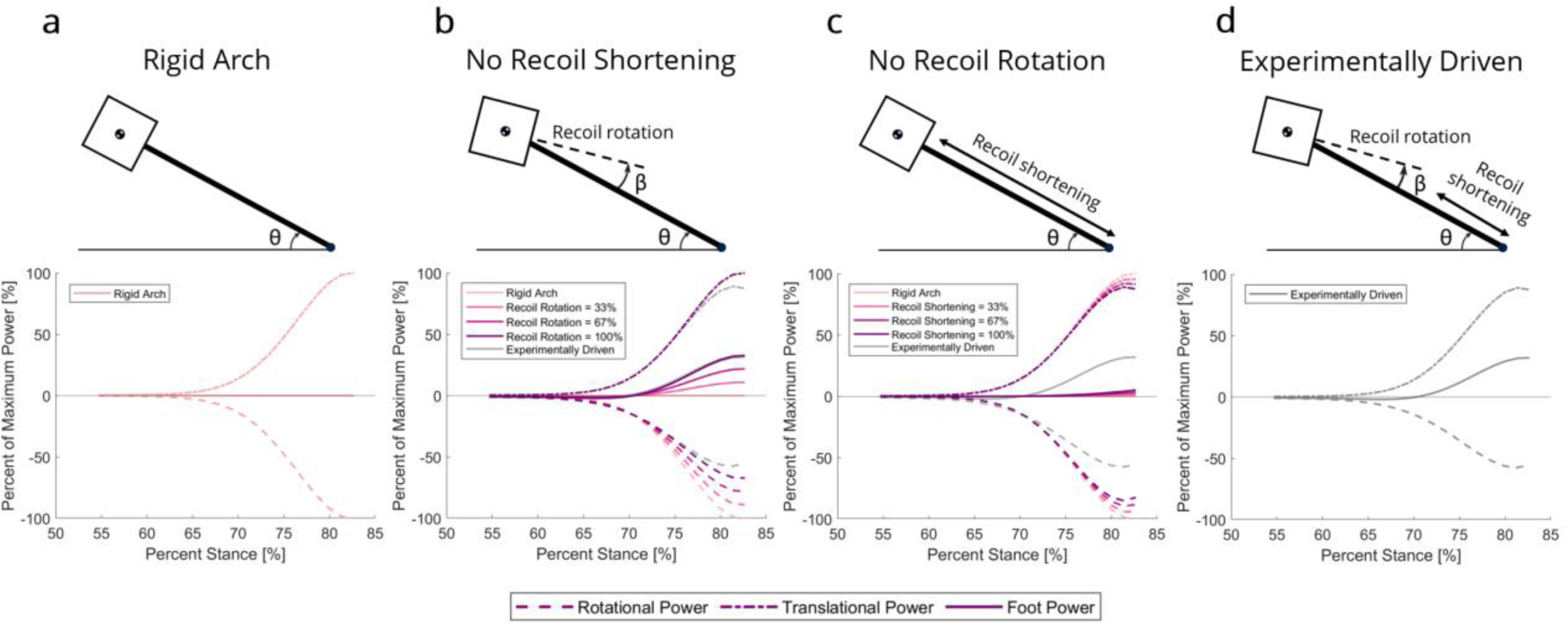
The net foot power (solid line), translational power (dash-dot line), and rotational power (dashed line) from the simple mathematical model for the (a) rigid arch case, (b) the no recoil shortening case, (c) the no recoil rotation case, and (d) the experimentally driven case, with the experimentally driven case shown with grey lines on the restricted shortening and rotation cases. For the no shortening (b) and no rotation cases (c), the nonzero variable was increased from 0% to 100% of the experimental values in increments of 33% represented by darkening lines.

**Figure 6:**
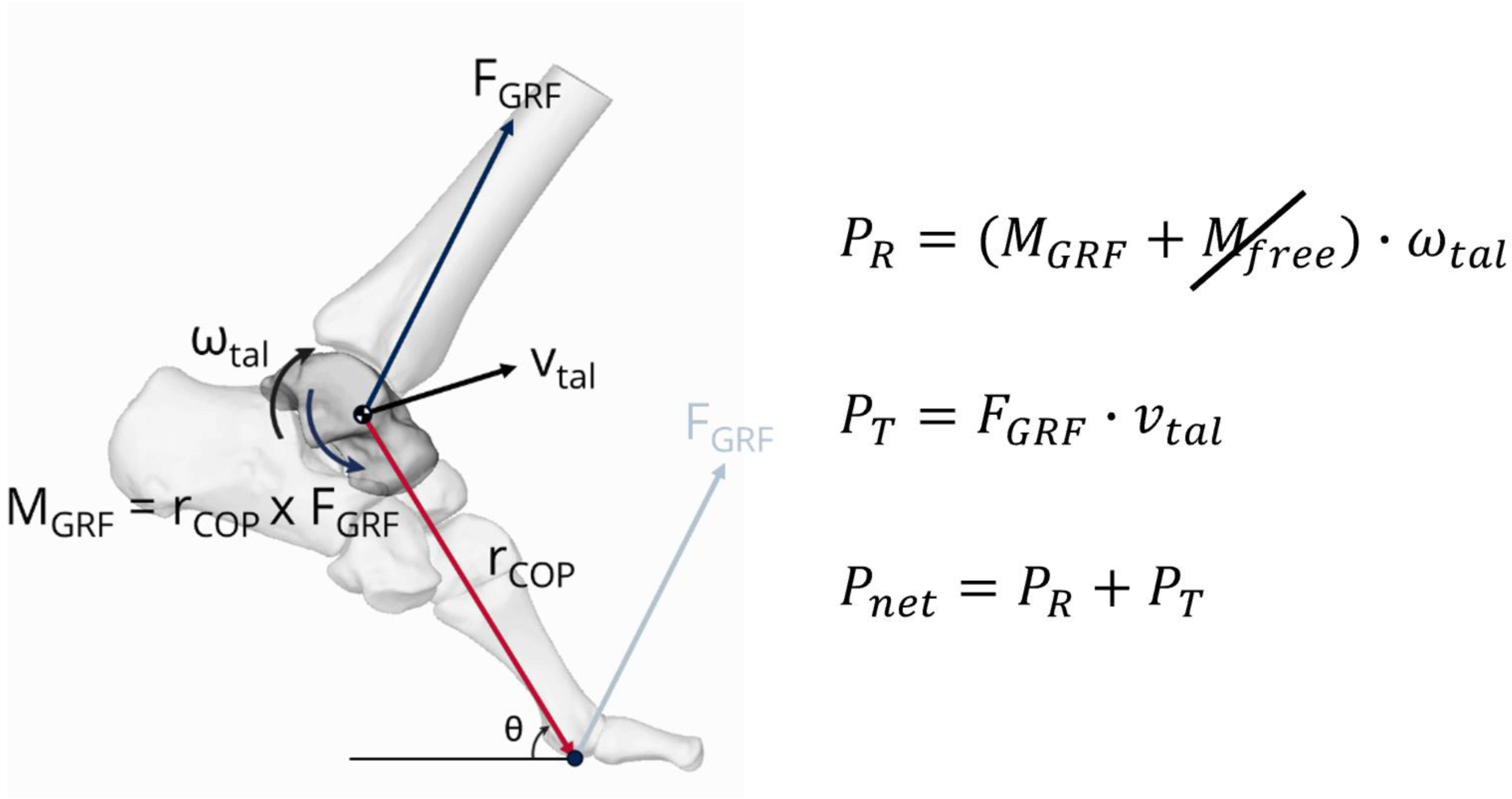
The moments and forces acting on the talus during propulsion and the rotational, translational, and net UD power equations. Since the model only accounts for sagittal plane motion, the free moment is zero.

We then set recoil shortening to zero but incrementally introduced recoil rotation relative to the rigid case (Figure 5b). Here, we see similar net power to the experimentally driven case despite not allowing the arch to shorten. Relative to the rigid arch case, gradually increasing posterior tilt of the talus decreased its global anterior tilt and reduced the magnitude of the negative rotational power, which resulted in increasing the net power until it nearly matched the experimental case. Posterior talus tilt did not affect the translational power.

When we set the recoil rotation of the talus to zero and introduced recoil shortening relative to the rigid case, we saw a large reduction in net power compared to the experimental case. Relative to the rigid case, increasing recoil shortening caused slight increases in net positive power (Figure 5c). This was due to the shortening lever arm causing the rotational and translational power to decrease, with rotational power decreasing slightly more than the translational power. A shorter lever simultaneously decreases the translational velocity of the talus (decreasing positive translational power) and decreases the moment caused by the ground reaction force a similar amount (decreasing the negative rotational power).

In summary, the simple model showed how the recoil rotation and the recoil shortening affected the foot power. We see that limiting linear propulsion of the talus (caused by recoil shortening) does not substantially impact the net foot power relative to the experimentally driven case. However, restricting the posterior tilt of the talus substantially decreases the net foot power. This is because the posterior tilt rotation of the talus drives the reduction of negative rotational power, allowing the net power to increase. This aligns with the prediction from the upright gait hypothesis for arch recoil^17^ – positive foot power acts to posteriorly tilt the talus in global, putting the talocrural surface in a more favourable position for gait, rather than directly propelling the body forwards and upwards.

### Timing of Foot and Ankle Power

The foot and talocrural joint start producing positive power simultaneously (overlapping confidence intervals). This occurs at the instant in stance that the arch is maximally flattened, the ankle is maximally dorsiflexed, and the ground reaction force on the forefoot is maximal (Figure 7). Positive foot power occurs at heel off for running, but after heel off for walking, since the ankle is still dorsiflexing when the heel lifts.

**Figure 7:**
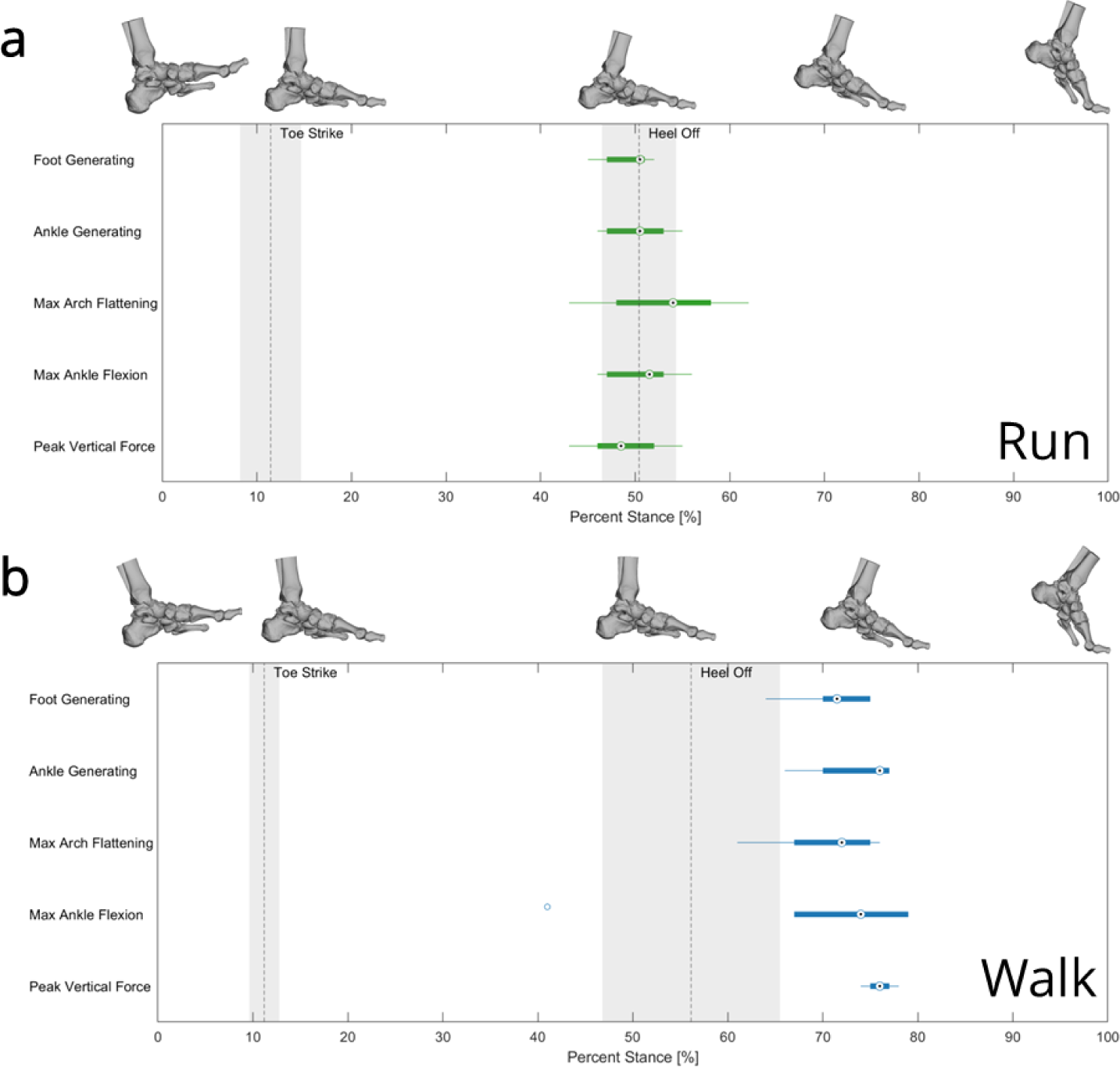
The timing of key events during stance for running (a) and walking (b). The key events are when the foot starts producing positive power (foot generating), when the ankle starts producing positive power (ankle generating), the point of maximum arch flattening, the point of maximum ankle dorsiflexion, and the point where the peak vertical ground reaction force under the forefoot occurs. The circles represent the median with boxes representing the 25^th^ to 75^th^ quartiles and bars representing the maximum and minimums. We see for both walking and running the key events all align. For running this occurs around the moment the heel lifts from the ground. For walking this occurs ∼20% of stance after the heel leaves the ground.

### Positive, Negative, and Net Work

The foot and talocrural work were distributed differently across walking and running (Figure 8). During running, the talocrural joint contributed substantially more positive work than the foot; however, the talocrural joint and foot contributed similar amounts of positive work during walking. The positive talocrural work generated more positive work during running compared to walking for all participants. The net talocrural work was larger in running compared to walking for 5 of 6 participants.

**Figure 8:**
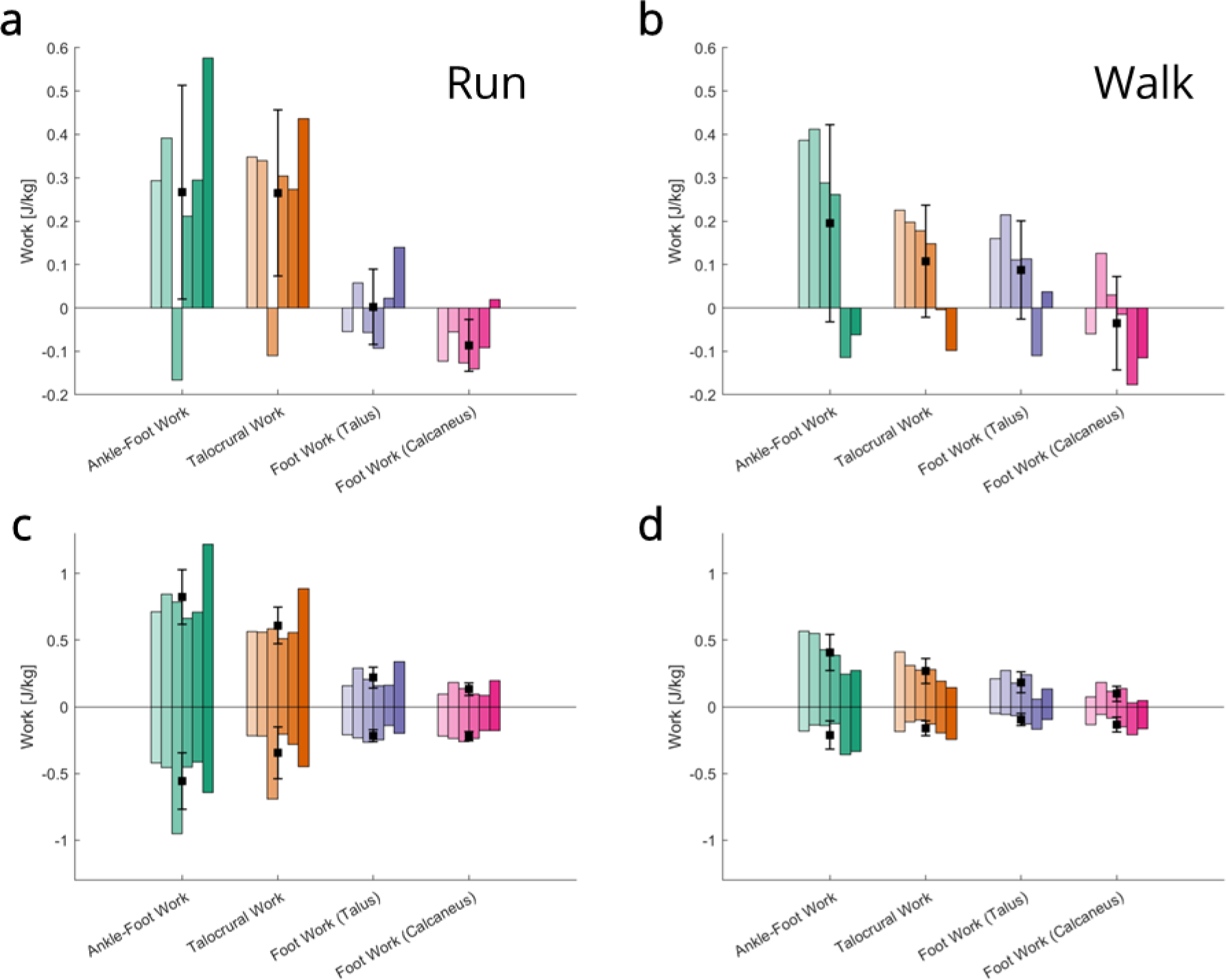
The net work for running (a) and walking (b) and the positive and negative work for running (c) and walking (d) produced over stance by the ankle-foot complex, the talocrural joint (ankle-foot minus foot), and the foot using the talus and the calcaneus as the proximal rigid segment. Each bar represents an individual participant with mean ± standard deviation shown by the black boxes with error bars.

The foot generated slightly more positive work during running compared to walking (4 of 6 participants), however the ranges overlapped substantially (running 0.16 to 0.34 J/kg, walking 0.05 to 0.27 J/kg). The foot did more negative work during running compared to walking for 5 of 6 participants (ranges of −0.14 to −0.26 J/kg running, −0.05 to −0.17 J/kg walking). This led to a small amount of positive (3 participants) or negative net work (3 participants) done by the foot for running, and positive net work done by the foot in walking (5 out of 6 participants).

There was a large variation among participants for the net work and positive/negative work across the foot, talocrural joint, and ankle-foot. Different participants had net positive and negative work done by the ankle-foot complex, talocrural joint, and foot (Figure 8).

### Foot Power with Talus Versus Calcaneus as Proximal Rigid Segment

Using the talus and calcaneus as the proximal rigid component produced different results for foot power, with the talus generating more power during propulsion (Figure 9). The net foot work was larger when using the talus, mainly because of the larger positive work (Figure 8).

**Figure 9:**
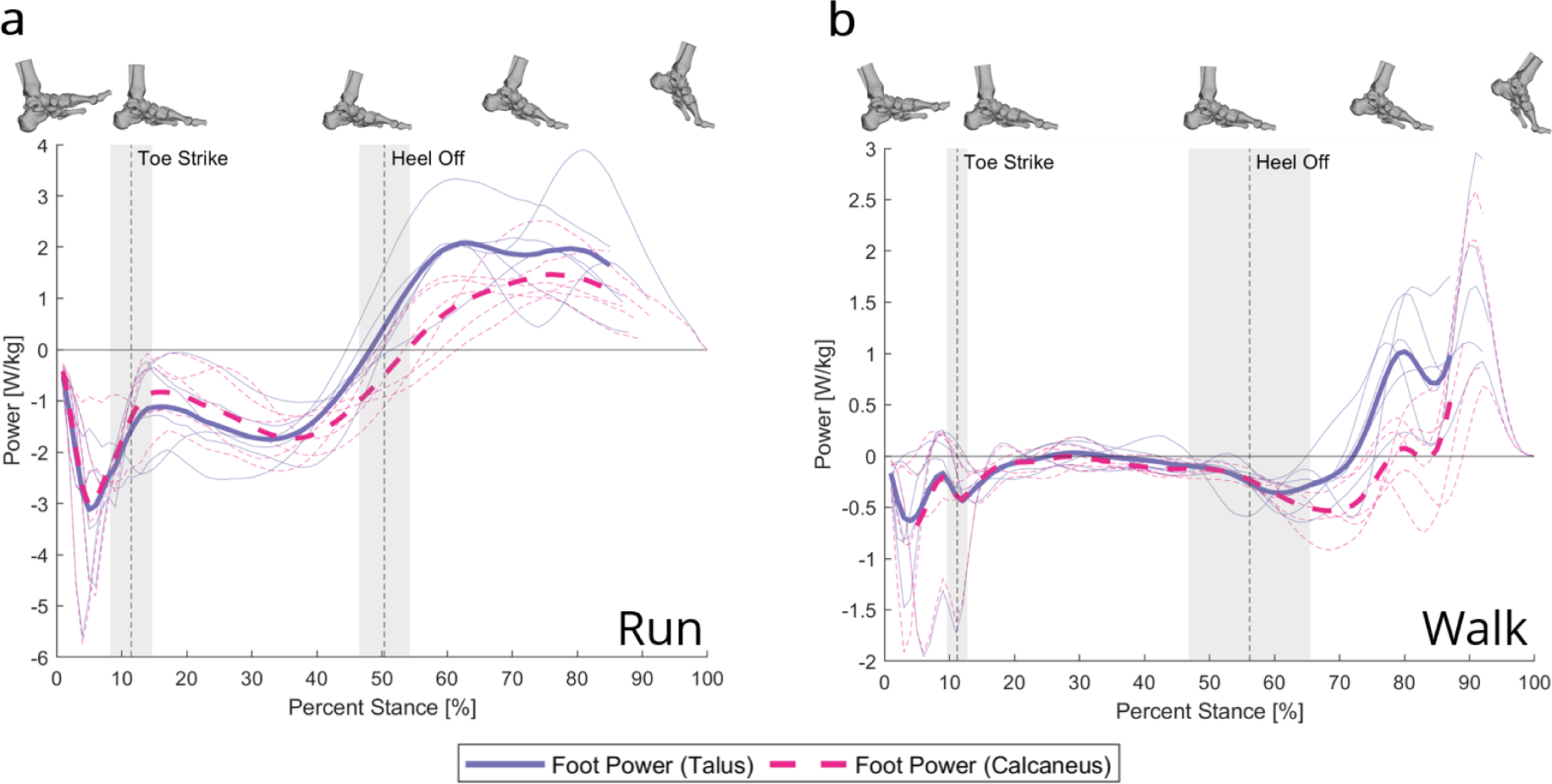
The unified deformable (UD) power over stance calculated using the talus and the calcaneus as the proximal rigid component for running (a) and walking (b). Thin lines represent each participant (n = 6), with the thick lines representing the mean (mean lines stop when data becomes unavailable for a single participant). Mean ± standard deviation of toe strike and heel off are indicated with dashed lines and gray shading.

## Discussion

This study used a novel method to measure foot and ankle power by combining the UD power model with highly accurate talus and tibia orientation and position measurements from BVR and kinetics from force plates. Using our simple mathematical model of the foot during propulsion, we found that net propulsive power occurs because the talus tilts more posteriorly during arch recoil than it would if the arch was rigid. Put differently, positive arch recoil power is used to slow down global talus anterior tilt caused by ankle plantar flexion. Our model shows that posterior talus tilt decreases the negative rotational power, which increases the net power. These findings are consistent with the arch recoil “upright gait hypothesis”.

The upright gait hypothesis for arch recoil has highlighted the importance of the kinematic relationship between the arch of the foot and the ankle for upright locomotion^17^. Talus posterior tilt from arch recoil keeps the talocrural surface more level, allowing the shank to remain upright and prolonging ground contact time^17^. As predicted by the upright gait hypothesis for arch recoil, we saw the power generated by the foot during propulsion goes into putting the talus into this more favourable position for push off. Using our simple model of the foot, we were able to independently restrict the linear actuation of the talus and the rotation of the talus caused by arch recoil to see how it affected net foot power. When the linear recoil of the arch was restricted, we saw similar power production from the deformable structures, indicating that, in our limited sample, arch shortening/lengthening does not substantially contribute to foot power during propulsion. However, when the backwards rotation of the talus was restricted, foot power was substantially decreased indicating that positive foot power during propulsion is linked with this talus posterior tilt. This model also helps explain how positive power can act to slow the rotation of a body. The arch recoil causes the talus to rotate backwards with respect to its global rotation, reducing its net global anterior tilt velocity. This reduction in velocity reduces the magnitude of the negative rotational component of the power, increasing the net power.

The timing of foot motion and power suggests that the arch’s function is critically linked to ankle motion and power generation. The arch and the ankle begin to simultaneously produce positive power in sync with the maximum arch flattening and ankle dorsiflexion. This coupling is driven by talus posterior tilt. Without posterior talus tilt, the magnitude of the negative rotational power would not decrease, and the net foot power would stay negative for longer. These findings support the existence of a mechanism linking arch recoil and ankle plantarflexion and suggests that the timing of arch recoil and ankle plantarflexion is essential for power generation during propulsion. This is intriguing given that there is evidence of a mechanical connection between the Achilles tendon and plantar aponeurosis^23–28^. However, we cannot further investigate individual tissue behavior using these methods.

The ankle-foot net work was greater than zero for 5 of 6 participants for running and 4 of 6 participants for walking, highlighting the importance of the ankle-foot complex for adding power to the system for walking and running. We also saw positive net work done by the foot for 5 of 6 participants for walking and a small amount of positive or negative net work varying amongst participants done by the foot during running. These findings challenge the idea that the foot acts as a spring-damper system for constant speed walking^3–5,13^ and running^12,13^ because the foot added power to the system for the majority of participants. However, we observed a large inter-participant variation with different participants performing positive or negative work across the foot and ankle that could not be explained by the small variations in speed or acceleration. We speculate that this could be because of variation in the morphology of the foot among participants^29^, because of neuromuscular control strategies that divide the work between the lower limb joints differently^30^, or step to step variations as only one step was captured. Due to the participant-specific nature of our results, we suggest avoiding “covering law” statements^31^ that discuss overall foot behavior in terms of group averages.

The distribution of work between the ankle and foot differs between walking and running. The foot performs slightly more positive work for running compared to walking, albeit with substantial overlap in the ranges. To deal with the larger positive ankle-foot work required for running, the ankle must contribute more positive work. This suggests muscular demands for faster/more dynamic locomotion are taken up by the ankle and that the energy contributions of the foot may be limited. During running, the negative work done during weight acceptance increases when compared to walking. This aligns with previous findings that found negative work done by the foot increased as running speed increased while positive work remained constant^12^ and suggests this relationship also extends to walking.

Studies measuring the UD power of the ankle-foot complex for walking and running are uncommon and our ankle-foot work values differed slightly from existing data. Takahashi et al.^3^ measured the ankle-foot power for walking and found an average of −0.010 ± 0.054 J/kg of net work done by the ankle-foot complex compared to our value 0.27 ± 0.25 J/kg. However, this was done on a set of pediatric participants and both datasets have large standard deviations. Although our average falls outside of the first standard deviation, we believe our results are aligned with this study. Previous studies^11,12^ and a preliminary analysis on an additional dataset (Supplemental S2) show that net work is closely related to speed. The pediatric sample walked at 1.30 ± 0.16 m/s while our sample walked at 1.58 ± 0.14 m/s. Both of these results matched a trendline fit to the speed and net work of the additional dataset (Supplemental S2). Also, as there is a large variation in both datasets there is a possibility that our sample just contained a higher percentage of “high power” walkers. We also calculated the UD power for the ankle-foot complex using the marker data and found that the results closely matched the power calculated using the BVR data (Supplemental S3).

There have been no studies applying the UD power model to the talus to measure foot power, but our foot power measured using the calcaneus was similar to previous studies for walking and running. In the same study as above, Takahashi et al.^3^ found the foot did −0.08 ± 0.02 J/kg of net work for walking compared to our value of −0.035 ± 0.11 J/kg when using the calcaneus as the proximal rigid segment. As before, we believe these values are aligned. For running, Kelly et al.^12^ measured the foot power using the calcaneus at three specific speeds and found a net work of −0.07 ± 0.08 J/kg at 2.2 m/s and −0.16 ± 0.10 J/kg at 3.3 m/s compared to our −0.09 ± 0.06 J/kg at 2.87 ± 0.09 m/s.

The foot power differed when measured using the talus as the proximal rigid segment versus the calcaneus for walking and running. We propose that the talus is an ideal choice for the proximal rigid segment when examining foot power for this specific question. The talus captures the entire foot as the distal, deformable structure. When used in tandem with the tibia as the proximal rigid segment, this novel application of the UD power model allows us to measure talocrural power by subtracting the foot power (talus as proximal rigid segment) from the ankle-foot power (tibia as proximal rigid segment). To our knowledge, the talus has not been used as the proximal rigid segment to capture foot power as it is inaccessible with traditional motion capture techniques; however, biplanar videoradiography (BVR) is able to capture talus motion, *in vivo*, during walking and running. We acknowledge that it is difficult to measure the position and orientation of the talus without BVR, which has its downsides in small measurement volume, long processing times, and small sample sizes. Because of this, it is important to understand the differences between the two methods. We found the foot power when using the talus had a similar positive translational power component but a smaller negative rotational power component during propulsion when compared to using the calcaneus. This meant that, when using the talus, the foot began generating positive power earlier in stance and had a larger peak power. Overall, the differences between the foot power profiles when using the talus versus the calcaneus suggests that the structures proximal to the calcaneus are important for positive power generation during propulsion for both walking and running.

This study has a few limitations. The UD power analysis is limited in its ability to distinguish what structures are generating/dissipating the power within a segment. Although previous studies have shown the arch stores and returns much of the power in the foot^3,6,8,10^, we can’t partition the foot power between the arch, intrinsic foot muscles, plantar fat pads, or other structures. Due to the long processing time for BVR studies, the sample size is only 6. However, BVR allows for highly accurate position and orientation measurements of the individual bones in the foot, including the talus which cannot be measured with optical motion capture. Also, one of the participants had tantalum beads implanted in their foot bones, giving highly accurate position and orientation measurements (0.1 mm translation, 0.1 degrees rotation). The results from this participant were consistent with the other 5, giving us confidence that the manually tracked data is accurate. An additional challenge with BVR is the small size of the collection volume, as bones in the hindfoot often leave the volume before the end of stance. Because of this, the power curves for the last 10%-20% of stance had to be extrapolated for 5 of the 6 participants for running and 4 of 6 participants for walking. These curves were manually inspected and closely matched previous data and ankle-foot work values closely matched values calculated using the optical motion capture data. BVR data collected with split force plates also requires a very specific foot placement by the participant. Participants were allowed as much time as required to practice their foot placement and were instructed to adjust their foot placement by changing their starting location, not deliberately targeting the volume.

## Conclusion

Using a combination of experimental measurements and a simple mathematical model we showed that the positive power generated by the foot during propulsion acts to rotate the talus backwards, keeping the talocrural surface level across both walking and running. This aligns with the upright gait hypothesis for arch recoil which suggests the arch recoils to keep the talocrural surface level, allowing us to walk and run upright. Additionally, we saw a kinetic link between the ankle and foot, further supporting the importance of their coordination during locomotion. These results update our understanding of the purpose of positive foot power during propulsion and support the importance of mobility of the arch for foot function. This has implications for the design of robots, prosthetics, and footwear, and could lead to innovation in clinical treatments of foot disorders that limit the mobility of the arch and ankle.

## Methods

### Experimental Protocol

5 healthy, physically active, participants (4F, 1M; mean ± s.d.: weight 67.4 ± 8.0 kg, height 1.70 ± 0.08 m, age 24.0 ± 3.1 years, 1 FFS, 4 RFS) participated in the study after IRB approval and informed consent. Retro-reflective markers were attached to the lateral and medial malleolus and femoral epicondyles. A cluster of 4 retro-reflective markers was fixed to the shank. Participants walked and ran over level ground at self-selected speeds (walking 1.58 ± 0.14 m/s, running 2.87 ± 0.09 m/s) in flexible, thin-soled, minimal shoes, (7.5 mm sole, 0 mm heel-toe drop, Xero Prio Shoes, Broomfield, CO, USA) while biplanar videoradiography (BVR) captured their foot bone motion (Skeletal Observation Lab, Queen’s University, Kingston, CAN) (walking 125 Hz, running 250 Hz) and an optical motion capture system (Qualysis Track Manager, Qualysis AB, Gothenburg, SWE) simultaneously recorded the 3D positions of the markers (walking 125 Hz, running 250 Hz). Participants landed on a split force plate set up (Force Plates, Location) (1000 Hz). Participants were given enough time to familiarize themselves with the experimental setup and to select a starting position, so their right foot landed in the BVR volume each time.

Additionally, one participant (M, weight 83 kg, height, 1.75 m, age 51, RFS) with tantalum beads previously implanted in their foot bones performed the same experimental protocol (Skeletal Observation Lab, Queen’s University, Kingston, CAN). These beads allow very accurate tracking of the motion of the foot bones and the data is considered a gold standard for BVR. The experimental protocol was approved by Queen’s University Health Sciences and Affiliated Teaching Hospitals Research Ethics Board. All participants gave informed consent prior to participation in the data collection.

A computed tomography (CT) scan was taken of each participant’s right foot while prone, with a maximally plantarflexed ankle (Revolution HD; General Electric Medical Systems, Chicago, IL, USA; resolution: 0.317 mm x 0.317 mm x 0.625 mm). The tibia, talus, calcaneus, and first metatarsal were segmented (Mimics, Materialise, Leuven, Belgium) and tessellated meshes were created using the masks. Coordinate systems of the bones were generated using the principal directions of the moment of inertia tensor of the meshes^32^ and we oriented at the centroid of each bone.

The orientation and translation of the tibia, talus, calcaneus, and first metatarsal while the foot was in the BVR volume was found using a previously described pipeline^33^. Briefly, the high-speed cameras were calibrated using a custom calibration object and the images were undistorted, using X-ray-specific software (XMALab, Brown University, USA). Partial volumes from the bone masks formed digitally reconstructed radiographs that were manually aligned with the two, undistorted x-ray views in custom software (Autoscoper, Brown University, Providence, RI, USA). A particle swarm optimization optimized the cross-correlation values. For the participant with implanted beads, the beads were tracked semi-manually in the two views (XMALab, Brown University, USA)^34^.

### Data Processing

The force data was down sampled to align with the sample rate of the optical motion capture and the BVR data. The quaternions of the BVR transformation matrices were filtered using a fourth order low-pass Butterworth filter with cut-off frequency of 12 Hz. For the beaded participant, the positions of the beads in the two 2D images were filtered with a cut-off frequency of 12 Hz.

Gait events were defined using force plate data with a threshold of 30 N. The impulse of anterior-posterior ground reaction force was found using trapezoidal integration to ensure participants weren’t accelerating or decelerating through the volume. The centre of mass (COM) of the pelvis segment was used to calculate the average speed through the volume.

As the entirety of the shank could not be captured using BVR, the position of the COM of the shank was found using optical motion capture and driven using BVR. A truncated cone was generated using the malleolus and epicondyle markers from a static optical motion capture trial (Visual3d, C-Motion, Germantown, USA). The COM of this cone was found relative to the cluster placed on the shank (T_1_). For each trial, the average position of the cluster (tracked with optical motion capture) relative to the COM of the partial tibia (tracked with BVR) during stance was found (T_2_). Using these two transforms, the position of the COM of the shank relative to the COM of the partial tibia was found (T_1_·T_2_). This allows the COM of the shank to be driven by the highly accurate transformation matrices of the tibia.

Joint angles were defined using Euler angles with a ZYX sequence where rotation about the x-axis was plantar/dorsiflexion, rotation about the y-axis was in/eversion and rotation about the z-axis was internal/external rotation (+/−). Arch deformation was defined as the x-axis rotation of the first metatarsal relative to the calcaneus. Ankle flexion was defined as the x-axis rotation of the talus relative to the tibia.

### Power and Work Analysis

We applied a unified deformable (UD) segment analysis to quantify the instantaneous power of the foot and the ankle-foot complexes. This method calculates the power of all structures distal to a segment by assuming all structures distal to the proximal rigid segment are massless and deformable^18^. We applied the UD power model to three different proximal rigid segments: the tibia to quantify the power produced by the ankle-foot complex, and the talus and calcaneus to quantify the power produced by the foot. The power of the talocrural (ankle) joint was calculated by subtracting the foot power (calculated using the talus) from the ankle-foot power (calculated using the tibia). The UD power is given by:

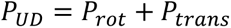

Where *P*_*rot*_ is the rotational component of power representing the distal structures rotating the proximal segment with moments and is given by:

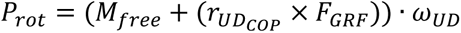

*P*_*trans*_ is the translational component of power representing the distal structures translating the proximal segment with forces and is given by:

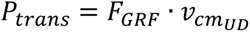

Where *F*_*GRF*_ is the ground reaction force, *M*_*free*_ is the free moment, and *ω*_*UD*_ is the angular velocity of the proximal rigid segment, *v*_*cm*_*UD*__ is the velocity of the COM of the proximal rigid segment and *r*_*UD*_*COP*__ is the displacement of the centre of pressure from the COM of the proximal rigid segment. Although this is a different derivation than originally given by Takahashi et al.^18^, it gives the same resultant UD power.

Work was calculated using trapezoidal integration of the instantaneous power data. Positive work and negative work were calculated by integrating over the parts of stance where power was greater than zero and less than zero, respectively. If the power curves ended before the end of stance due to a bone leaving the BVR volume, the curves were extrapolated using cubic splines such that the power was zero at toe off. All curves were manually inspected to ensure consistency with previous studies.

### Simple Foot Model

The UD power for the two-dimensional simple foot model was calculated using basic principles of motion (for additional details see Supplemental S1). Briefly, the velocity and acceleration of the mass (given mass of 1 kg) was found by taking the first and second derivative of a vector from the pivot point to the mass. The reaction force at the pivot was found using Newton’s Second Law. The UD power was then calculated with the ground reaction force as the reaction force at the pivot, the centre of pressure being the pivot point, and the *ω*_*UD*_ being the rotation of the lever about the pivot point plus the rotation of the mass with respect to the lever. The model had input parameters of the angular acceleration of the mass with respect to the lever, the angular acceleration of the lever about the pivot point, and the rate of change of the shortening velocity of the lever. Euler’s method was used to solve for lower order values at each time step given initial values. To limit the recoil rotation, the angular acceleration of the mass with respect to the lever was scaled by factors of 0, 0.33, and 0.67. To limit the recoil shortening the rate of change of the shortening velocity of the lever was scaled by factors of 0, 0.33, and 0.67. The angular acceleration of the lever about the pivot point was unchanged across all conditions.

