## Supplemental for "Reassessing the Role of Foot Power in Human Gait"

### S1 – Simple Model Equations

The simple model was driven by Newtonian mechanics. The model was composed of a point mass that was able to rotate on the end of a massless lever (Figure S1). This was fixed to the ground at a pivot point. The levering of the system about the pivot point represented the foot levering about the metatarsal heads during ankle plantar flexion. The mass tilting backwards represented the posterior tilt of the talus caused by the recoil of the longitudinal arch. Position, velocity, and acceleration of the mass relative to the fixed pivot point was found by getting the position of the mass relative to the fixed point in inertial frame $\hat{\imath} - \hat{\jmath}$ then taking the vector derivative.  Newton's second law was used to find the reaction force at the pin joint (which represented the ground reaction force). The unified deformable (UD) power of the system was then calculated using the UD power equations.

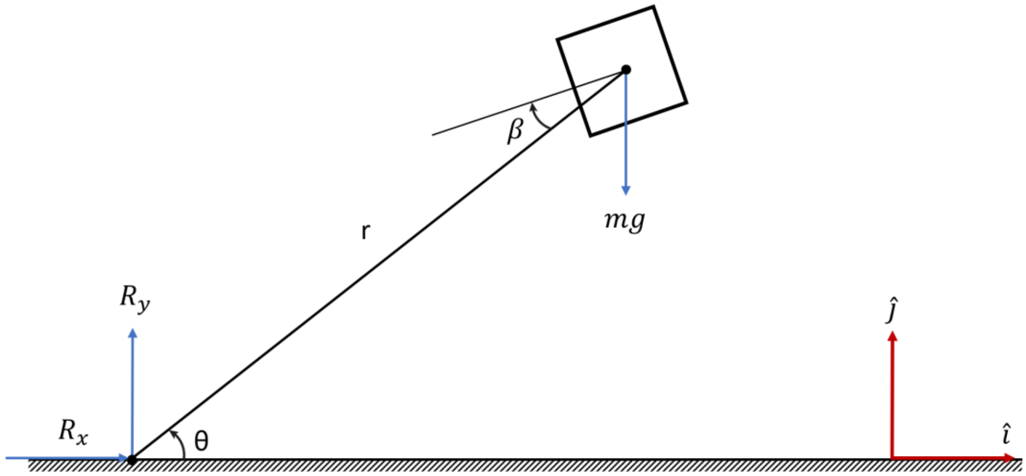


*Figure S1: Diagram of the simple 2D model showing the mass on a massless lever arm that pivots about a pin joint. The force of gravity on the mass and the reaction force at the pin is shown in blue. The inertial reference frame is shown in red.*

Position, velocity, and acceleration of mass relative to pivot point:

$$\overrightarrow{r_m} = rcos(\theta)\hat{\imath} + rsin(\theta)\hat{\jmath}$$

$$\overrightarrow{v_m} = \left(\dot{r}\cos(\theta) - rsin(\theta)\dot{\theta}\right)\hat{\imath} + \left(\dot{r}\sin(\theta) + rcos(\theta)\dot{\theta}\right)\hat{\jmath}$$

$$\overrightarrow{a_m} = \left(\ddot{r}\cos(\theta) - 2\dot{r}\sin(\theta)\,\dot{\theta} - rcos(\theta)\dot{\theta}^2 - rsin(\theta)\ddot{\theta}\right)\hat{\imath}$$

$$+\left(\ddot{r}\sin(\theta) + 2\dot{r}\cos(\theta)\,\dot{\theta} - rsin(\theta)\dot{\theta}^2 + rcos(\theta)\ddot{\theta}\right)\hat{\jmath}$$

Force balance:

$$\sum F_i = R_x = ma_x$$

$$R_x = m\left(\ddot{r}\cos(\theta) - 2\dot{r}\sin(\theta)\,\dot{\theta} - rcos(\theta)\dot{\theta}^2 - rsin(\theta)\ddot{\theta}\right)$$

$$\sum F_j = R_y - mg = ma_y$$

$$R_y = m\left(g + \ddot{r}\sin(\theta) + 2\dot{r}\cos(\theta)\,\dot{\theta} - rsin(\theta)\dot{\theta}^2 + rcos(\theta)\ddot{\theta}\right)$$

Power equations:

$$P_{trans} = \vec{F}\cdot\vec{v} = \vec{R}\cdot\overrightarrow{v_m}$$

$$P_{trans} = m\left(\dot{r}\cos(\theta) - rsin(\theta)\dot{\theta}\right)\left(\ddot{r}\cos(\theta) - 2\dot{r}\sin(\theta)\,\dot{\theta} - rcos(\theta)\dot{\theta}^2 - rsin(\theta)\ddot{\theta}\right)$$

$$+\,m\left(\dot{r}\sin(\theta) + rcos(\theta)\dot{\theta}\right)\left(g + \ddot{r}\sin(\theta) + 2\dot{r}\cos(\theta)\,\dot{\theta} - rsin(\theta)\dot{\theta}^2 + rcos(\theta)\ddot{\theta}\right)$$

$$P_{trans} = m\left(\dot{r}r\dot{\theta}^2 + \ddot{r}\dot{r} + r^2\dot{\theta}\ddot{\theta} + g\dot{r}\sin(\theta) + gr\dot{\theta}\cos(\theta)\right)$$

$$P_{rot} = \left(\vec{r}\times\vec{F} + \cancel{\overrightarrow{M_{free}}}\right)\cdot\vec{\omega} = \left(\overrightarrow{r_m}\times\vec{R}\right)\cdot\left(\dot{\theta} + \dot{\beta}\right)$$

$$P_{rot} = -m\left(\dot{\theta} + \dot{\beta}\right)\Big(rsin(\theta)\left(rcos(\theta)\dot{\theta}^2 + 2\dot{r}\sin(\theta)\,\dot{\theta} - \ddot{r}\cos(\theta) + rsin(\theta)\ddot{\theta}\right) +$$

$$rcos(\theta)\left(-rsin(\theta)\dot{\theta}^2 + 2\dot{r}\cos(\theta)\,\dot{\theta} + \ddot{r}\sin(\theta) + rcos(\theta)\ddot{\theta} + g\right)\Big)$$

$$P_{rot} = -m\left(\dot{\theta} + \dot{\beta}\right)\Big(2\dot{r}\dot{\theta} + r^2\ddot{\theta} + rgcos(\theta)\Big)$$

For completely rigid arch ($\dot{r} = 0, \ddot{r} = 0, \dot{\beta} = 0$):

$$P_{rot} = -m\dot{\theta}\left(rcos(\theta)g + r^2\ddot{\theta}\right)$$

$$P_{trans} = m\dot{\theta}\left(rcos(\theta)g + r^2\ddot{\theta}\right)$$

$$\Longrightarrow P_{trans} = -P_{rot}$$

### S2 – Net Work vs Speed from Additional Dataset

An additional dataset of 7 participants walking at slow, medium, and fast speeds and running (all self-selected) was used to see how speed could influence the ankle-foot work. The participants walked and ran over force plates while optical motion capture (Qualysis Track Manager, Qualysis AB, Gothenburg, SWE) recorded the 3D positions of a cluster of 4 markers fixed to the shank and markers fixed to the lateral and medial malleolus and femur epicondyles (shank segment) and left and right anterior and posterior iliac spine (pelvis segment) for 2-3 cycles of gait. Each cycle was analyzed independently. The centre of the four markers fixed to the pelvis was used to calculate the velocity of the centre of mass. The ankle-foot power was calculated using Visual3d (C-Motion, Germantown, USA). The ankle-foot work was calculated using trapezoidal integration over stance. A linear model was fit to the data using MATLAB's (Mathworks, Natick, USA) fitlm function.

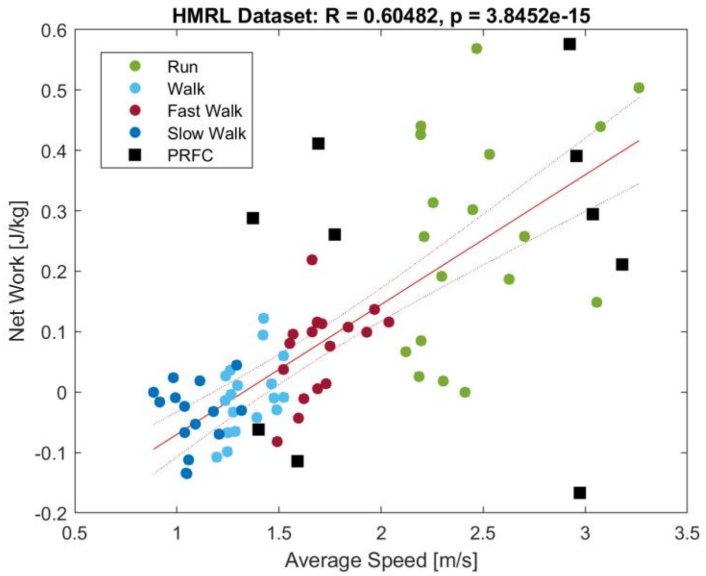


*Figure S2: The net ankle-foot work plotted versus speed for the additionally dataset with linear model fit. The dataset analyzed in the study is shown by the black squares.*

### S3 – Ankle-foot Power Calculated from Markers

To understand how calculating ankle-foot power using the biplanar transforms differed from the traditional optical motion capture techniques, we calculated the net work done by the ankle-foot complex using the marker trajectories for three participants (50% of the dataset). We saw good alignment between the two methods showing that using the tibia transforms was a valid way to calculate the ankle-foot power using the unified deformable model.

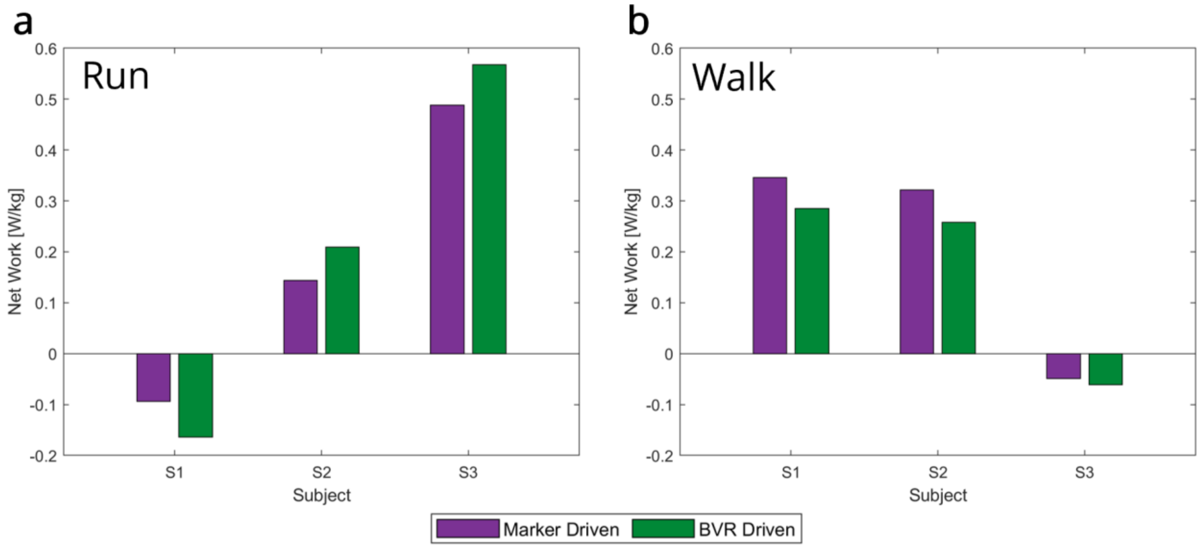


*Figure S3: The net ankle-foot work calculated using optical motion capture and biplanar videoradiography.*
